# EGIDExpress: An Interactive Shiny Web App to Visualize and Share Large Biological Datasets

**DOI:** 10.1101/2022.10.03.510709

**Authors:** John A. Besse, Garrett A. Osswald, Adina Y. Ballaban, Julie M. Caldwell, Marc E. Rothenberg

## Abstract

Biomedical research on rare diseases faces challenges such as low availability of biological specimens to study and limited funding. Eosinophilic gastrointestinal diseases (EGIDs) are rare conditions associated with inappropriate inflammation and the accumulation of eosinophils in various segments of the gastrointestinal tract. We aimed to build a repository of large datasets related to EGIDs that would be easily browsable, interpretable, and accessible in order to facilitate data sharing and hypothesis generation. Using the R-code based package Shiny, we built a website that allows visualization of multiple types of datasets including microarray, RNAseq, protein array, single-cell RNAseq, and ChIPseq. Users can access EGIDExpress (https://egidexpress.research.cchmc.org/) to browse data on a per-gene basis and to generate graphic representation of the data. Additionally, users can download the processed data to initiate their own analyses. Within 34 months of launching EGIDExpress, over 2400 users from 37 countries and 37 states within the United States accessed the site. Overall, EGIDExpress is accelerating research on EGIDs and provides a prototypic platform for broad research data sharing.

## Introduction

Eosinophilic gastrointestinal diseases (EGIDs) are rare allergic diseases that affect various segments of the gastrointestinal tract with eosinophil-predominant inflammation, resulting in physiological dysfunction of the organs, a variety of clinical symptoms, and decreased quality of life ^1^. Although diet and off-label anti-inflammatory treatments improve symptoms and histology, only one FDA-approved drug (dupilumab) exists for any of these diseases ^2, 3^. Moreover, current therapies are not curative; once any treatment is withdrawn, the diseases relapse. These factors underscore the need for a better understanding of the molecular and cellular processes underlying these diseases so that additional targets for treatment or prevention of diseases can be identified.

The generation of big data sets over the past 20 years has driven basic science and translational research to accelerate. These large data sets, however, require a substantial investment of time and financial capital to not only produce but also to maintain. In the case of rare diseases such as EGID, the most valuable samples of interest are often scarce and not readily accessible or shareable once they are generated. Furthermore, the multiple different types of experimental data sets make cataloging and accessing the big datasets difficult. These factors are compounded by the frequent turnover of staff, wherein data sets and critical associated information tend to get lost upon departure of the person leading the acquisition of the data set. These factors make production of a platform that is easily amenable to adding, distributing, and visually sharing the data a challenging yet worthy pursuit. Having such a platform would allow unpublished data to be privately accessible among laboratory members and trusted collaborators, and published data could be made readily accessible to both the research community at large and the general public.

We aimed to produce a platform to address these issues using a web-based R shiny app. The main goal of the app was to quickly visualize data from different types of experiments gathered from various platforms, allowing researchers to generate testable hypotheses or support previous ones. The goal of the visual format was to break down data easily and quickly and make various aspects of the experimental design readily interpretable; for example, to present the data such that it is easy to derive the sample source and characteristics. The app covers two aspects, the first being the ability to observe multiple datasets each from a limited perspective (e.g., one can briefly yet quickly examine small parts of the entirety of the dataset by selecting a gene of interest and observing its behavior in multiple experiments). This aspect includes generation of statistical analysis between sample groups for the chosen gene. The second aspect is that it acts as a repository for data and metadata for further analysis. The app presents data through two separate user interfaces: one for published data that is publicly accessible and one for unpublished data that is password-protected. Through Google analytics, site traffic is monitored providing a means to foster collaborations among individuals identified by geographic location. Overall, EGIDExpress provides a model platform for compiling medical research datasets and making them readily browsable, interpretable, and shareable.

## Materials and Methods

EGIDExpress is a web app built with the Shiny R package implementing a user-friendly graphical interface that offers the ability to visualize data in several formats, to view the statistical significance of the results, view relevant sample information, and download raw data sets in CSV formatted files ^4^. Any dataset that can be formatted into an expression matrix, that is, a table of gene or protein expression value versus sample information, is compatible with EGIDExpress. The computational core consists of the functionalities of gridExtra_2.3, SeuratObject_4.0.1, Seurat_4.0.2, dashboardthemes_1.1.3, shinydashboard_0.7.1, shinyjs_2.0.0, ggplot2_3.3.3, shiny_1.6.0 R, Gviz_1.34.1, and ensembldb_2.14.1 packages^5, 6, 7, 8, 9^. The R standard library packages utilized include stats, graphics, grDevices, utils, datasets, methods, and base. The graphical portion of the app has been implemented through the functionalities of the ggplot2_3.3.3 package^10^. Statistics are computed from a built-in package (stats). Single cell data sets have been implemented through the functionalities of the SeuratObject_4.0.1 and Seurat_4.0.2 packages^11^. Graphical icons for buttons and headers are from the fontawesome_0.2.2 package. The data (expression matrices, Seurat objects, BAM files, or gene name vectors) and metadata for each dataset are stored in RData files as data frames. There are 3 RData files; one for the public site, one for the private site, and one for the ChIPseq site. All data can be downloaded in a Unix/Linux CSV file in an expression matrix format. Mobirise_5.6.5 was used to create the homepage. Google analytics collects usage data, and Google Data Studio displays the data. The app is hosted using an R shiny server (version 1.5.12.933) on an 8 GB RAM Linux system (Linux bmiegidexpresp1.chmcres.cchmc.org 3.10.0-1160.45.1.el7.x86_64 #1 SMP Wed Oct 13 17:20:51 UTC 2021 x86_64 x86_64 x86_64 GNU/Linux) that is maintained by the Cincinnati Children’s Hospital Division of Biomedical Informatics. EGIDExpress is available online (https://egidexpress.research.cchmc.org/). We tested website functionality on macOS, Windows, Android, and iOS systems. A brief YouTube video tutorial describing the main features of EGIDExpress can be found at the following URL: https://www.youtube.com/watch?v=8pzbU-L_N_8.

The R-shiny reactivity model has been used to obtain user input, retrieve data, and render it visually. It supports the large data volume and the responsiveness of the web app by using reactive functions to display small amounts of the data volume based on user input (e.g., dataset selection, gene selection, or graphical selection). For further optimization, all data are loaded from the respective RData file on startup to eliminate loading individual CSV or TXT files containing the expression matrix or sample information data for each dataset. Once the site completes its initial loading, navigating between various datasets, genes, and other options requires no further loading. Finally, the code used to generate EGIDExpress has been set up so that adding a new dataset requires minimal alteration to expedite the addition of new data sets or experiment types.

To protect unpublished datasets yet allow for their sharing among lab members, a separate password-protected version of EGIDExpress was created with the addition of the sodium_1.2.0 package^12^. The display of CHIP-seq data is also available within the password-protected site. Version control of EGIDExpress is managed using a local GIT repository and the gitcreds_0.1.1 package. This functionality provides update history and allows multiple developers to be working simultaneously. Individual changes can be merged before publication to the website.

### EGIDExpress main interface

The overall structure of user interaction with EGIDExpress is outlined in the user flow diagram in Fig 1. When a user visits the EGIDExpress website the user lands on the homepage that describes the app, has an instruction video, links to external websites, and shows the geographical location and quantity of recent site visitors. Upon the user pressing the Browse Data button, the main R shiny interface is opened (Fig 2).

**Figure 1.**
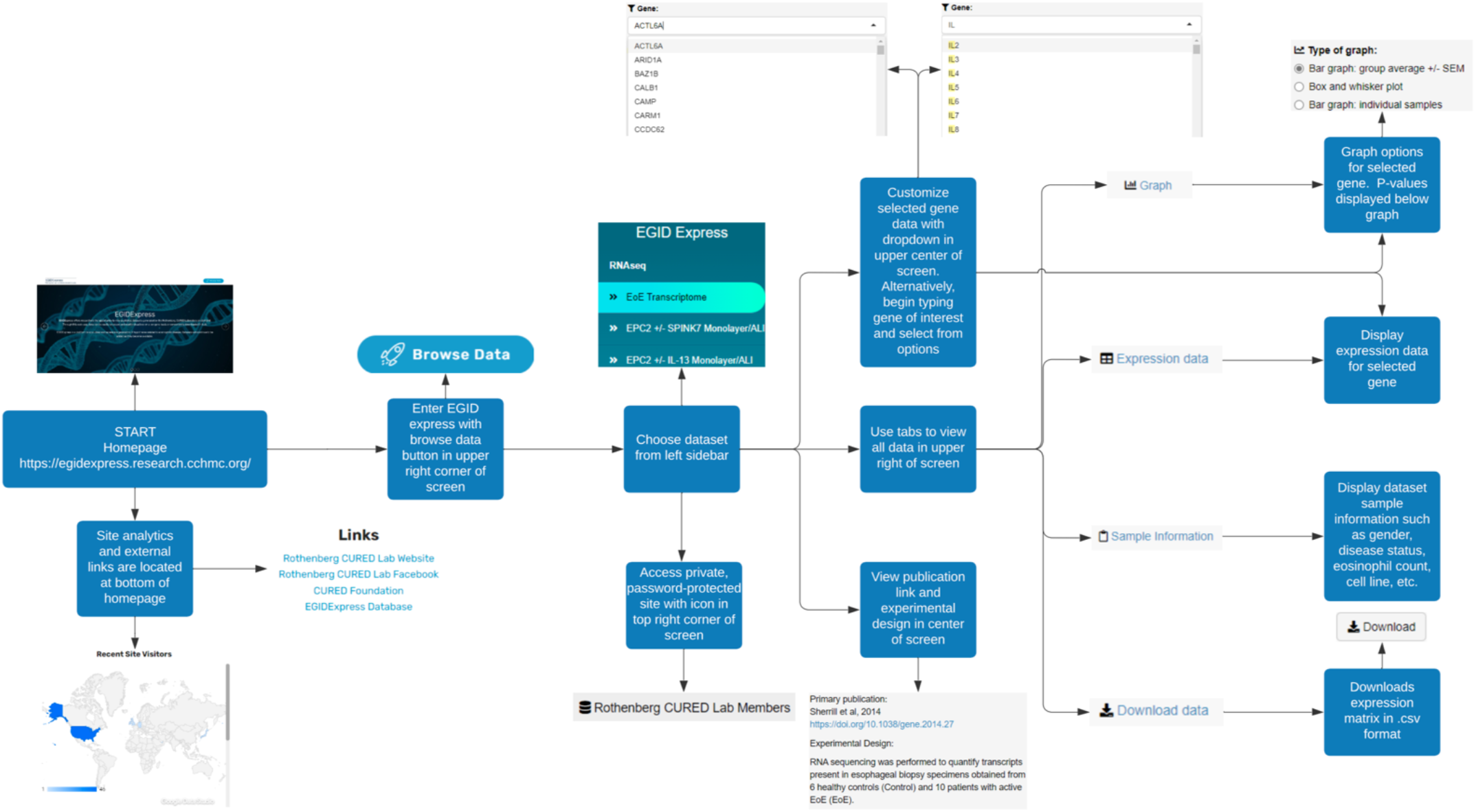
EGIDExpress organization. The primary purpose of EGIDExpress is to browse data from a given experiment on a per gene basis. The EGIDExpress homepage describes its main goal, has an instructional video, and displays site visitor locations. Upon entering the app, the first module of EGIDExpress allows the user to choose the experiment. This causes the second module of the app to render the gene list associated with the experiment and the third module of the app to render the static sample information and expression matrix associated with the experiment. The user then chooses a gene and graph output type. This then renders the data in graphical and text format, and processing of the data occurs to output statistical analysis of the data.

**Figure 2.**
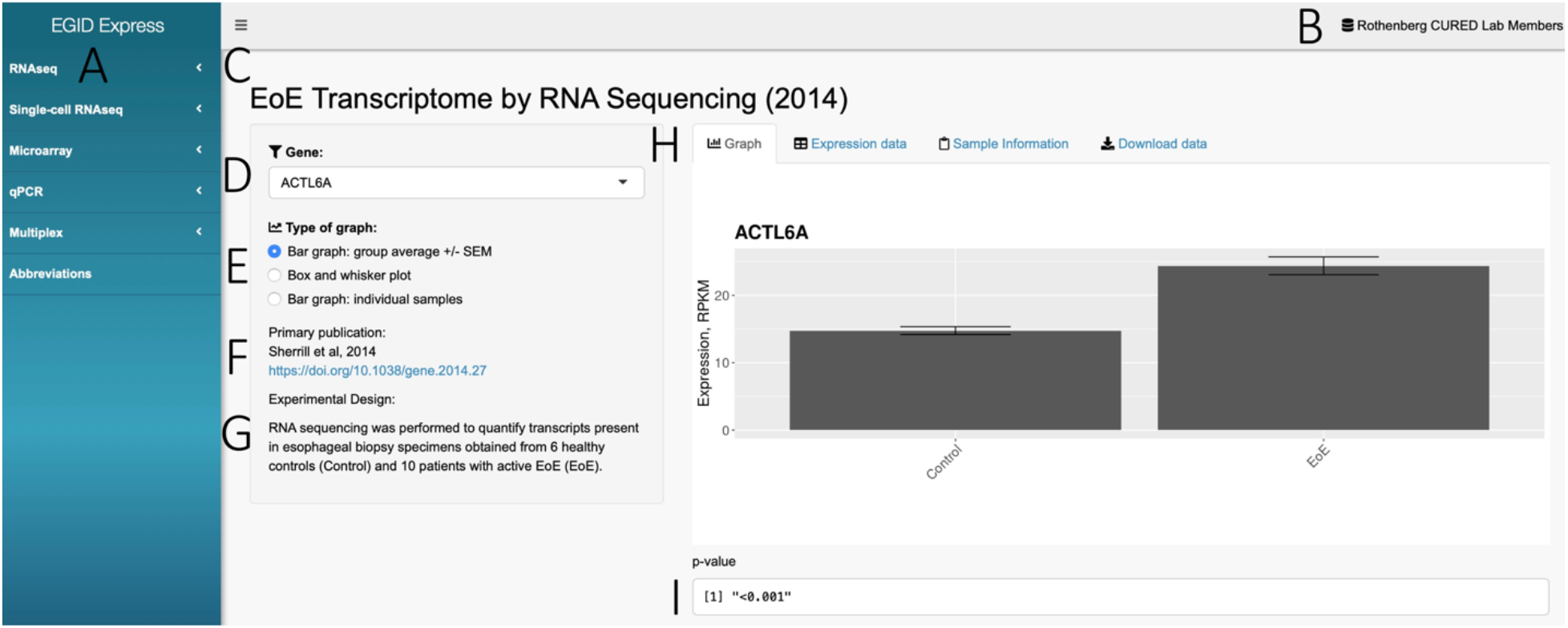
Example EGIDExpress page layout. (A) Sidebar with all experimental categories; choosing a category reveals a dropdown list of all relevant experiments. (B) Header button to access private version of EGIDexpress for unpublished data. (C) Dataset title with publication year. (D) Gene selection dropdown menu. (E) Type of graph selection. (F) Primary publication reference for dataset. (G) Brief experimental design description. (H) Tabs for viewing dataset graph, data, sample information, and downloading data. (I) Statistical analysis between experimental groups, if applicable.

First, the user must choose a platform and then a dataset of interest from the left sidebar (Fig 2A). Datasets are organized by the platform that generated the results, and they include next generation sequencing results (bulk RNAseq [RNAseq], single-cell RNA sequencing [Single-cell RNAseq], chromatin immunoprecipitation sequencing [ChIPseq, only available on password protected site]), microarray (Microarray), quantitative real-time PCR (qPCR), and multiplex protein assays (Multiplex). Selection of a platform reveals a dropdown list of all relevant experiments. Datasets are named by an abbreviated description of the biological specimen analyzed, and in some cases, the stimulatory molecule and/or growth conditions to which the biological specimens were subjected. Additionally, a list of all abbreviations used within the app can be accessed by clicking the Abbreviations button on the left sidebar. A header button in the upper right (Fig 2B) provides access to the password-protected version of EGIDExpress for Rothenberg CURED Lab members. A header button on the password-protected site further links to the ChIPseq site or back to the public site.

Upon choosing a dataset, a central window opens revealing a descriptive title of the experiment with the year of the primary publication noted (Fig 2B) and a box which displays a dropdown menu of genes (Fig 2C). Below the gene selection are the graphical display options (Fig 2E), the primary publication reporting the data with its manuscript URL (Fig 2F), and text describing the experimental design (Fig 2G). The user may select a gene from the dropdown menu. The genes are populated from the row names of the expression matrix. The dropdown menu has a display limit of 200 genes; to find any gene beyond that list, the user must begin typing the gene name, and the interface will search for and display any gene name that has the entered characters as a substring. Selecting a gene renders a graph of the dataset filtered for this gene. For all datasets except single-cell RNAseq and ChIPseq datasets, this selection also filters for the gene’s expression data and populates it within the expression data tab.

Below the dropdown menu, the graph type can be changed by selecting from a radio button list of graph types implemented for that dataset. For all current datasets excluding those obtained from the single-cell RNAseq platform or ChIPseq datasets, the choices are as follows: bar graph: group average +/-standard error of the mean (SEM); box and whisker plot; and either bar graph: individual samples or dot plot: individual samples. For single-cell RNAseq datasets, the choices for graphs are as follows: Feature Plot, Violin Plot, or Ridge Plot; the current single-cell RNAseq dataset additionally has the choice of whether to display the data grouped by subject disease state or to combine all subjects’ data. For ChIPseq datasets, the choice of graph is pre-set, but several display options must be specified. First, the number of base pairs 5-prime (before) and 3-prime (after) of the transcriptional start site (TSS) of the gene must be chosen (defaults to 5000 each) using the up-down controls. Display options include solid fill or outline only, no fill. There is also a choice to overlay all samples on one track. Color choice for the observation or types of chromatin modification (e.g., assay for transposase-accessible chromatin with high-throughput sequencing [ATAC-Seq], histone H3 lysine 27 acetylation [27Ac], histone H3 lysine 4 tri-methylation [Me3]) are made via dropdown menu.

To the right of the box containing the dropdown menu, graph choice, and experimental description is a box which contains choices for data output based on the choices specified in the left box (gene, graph type) as well as visualization of sample information and downloading data. Output choices are specified by tabs. A maximum of four tabs are available per dataset, namely Graph, Expression data, Sample information, and Download data (Fig 2H). The default is Graph; this represents the only choice for Single-Cell RNAseq datasets. The graph is generated based on the gene and type of graph specified in the left box. In all non-single-cell RNAseq data sets containing at least 2 samples per experimental group, statistics (p-value generated by Student’s T-test) are generated for comparison of groups in all possible combinations and can be visualized under the graph (Fig 2I). The graphs are generated as images by the program, and the user can interact with them using a right click to save them or enlarge them if viewed in a separate tab. If the desired output is numerical data on a per sample basis as opposed to visualization of the data in graphical format, the user can choose the Expression data tab. This tab lists the sample name vs. expression value (units specified per experiment) for each individual sample in a table format according to the gene chosen in the dropdown menu. The third and fourth tabs contain meta-information for the experiment at-large and do not depend on any user selection. To understand additional information about each sample, the user can browse the Sample Information tab. This tab lists the sample title, which corresponds to the title listed in the Graph and Expression data tabs, and specifies information associated with each such sample, including information about the biological specimen, culture conditions, treatments, number of replicates, disease activity, subject demographics, among others. Finally, if a user wishes to obtain the data for all genes in the dataset, the Download data tab can be accessed, and upon pressing the Download button, the expression matrix will be downloaded in Unix/Linux CSV format. The expression matrix is in a standard format of gene vs. experimental condition.

### EGIDExpress Outputs

Upon opening the main interface and after choosing a dataset to browse, the user has the option of viewing four tabs in the right frame on the webpage. The visualization options presented in the first tab depend on the type of dataset. Currently in EGIDExpress, there are 3 main categories of datasets relative to the graphical outputs: 1) those with data that can be expressed in the format of an expression matrix, e.g., RNAseq, microarray, qPCR, multiplex protein; 2) those that are expressed as a Seurat object in an RDS file (e.g., single-cell RNAseq); and 3) those that can be expressed as BAM files (e.g., ChIPseq).

The first category of datasets includes those that can be expressed in the format of a simple expression matrix. This matrix can be in table format with the sample on the x-axis and gene name on the y-axis and expression value in the grid. In most cases for gene expression experiments (RNAseq, microarray, qPCR), this represents per gene data that has been normalized to a housekeeping gene (microarray, qPCR) or per reads (RNAseq). The expression matrix format is also amenable to readouts that do not necessarily require normalization and simply have an absolute value for each analyte (e.g., protein multiplex).

Each dataset in this first category has the option to visualize the chosen gene in one of three graph types that are specified just under the gene dropdown menu and displayed in the Graph tab. The first option in all cases is Bar graph: group average +/-SEM. This renders a graph that shows the gene name at the top of the graph, the units of measure on the y-axis, and the experimental group titles on the x-axis (Fig. 3A). The bars always originate at 0 and the height represents the average value for the group. The error bars represent the standard error of the mean. The y-axis scale is scaled based on the defaults in the ggplot package which is used to generate and render the graphs. The second option in all cases is box and whisker plot. This renders a graph that shows the gene name at the top of the graph, the units of measure on the y-axis, and the experimental group titles on the x-axis (Fig. 3B). The y-axis scale is scaled based on the defaults in the ggplot package which is used to generate and render the graphs. The third option is either a bar graph or a dot plot of all individual samples. For those datasets where the option is a bar graph, this renders a graph that shows the gene name at the top of the graph, the units of measure on the y-axis, and the individual sample names on the x-axis (Fig. 3C). The bars are shaded by their main experimental group definition, which is indicated in a key to the right of the main graph. For the datasets where the option is a dot plot, this renders a graph that shows the gene name at the top of the graph, the units of measure on the y-axis, and the individual sample names on the x-axis (Fig. 3D). The dots are colored by their main experimental group definition, which is indicated in a key to the right of the main graph. We chose to use either bar graphs or dot plots based on the size of the dataset and for the best visual clarity of the graph. For both the bar and the dot graphs, the y-axis is scaled based on the defaults in the ggplot package which is used to generate and render the graphs. For each of the three options, the Student’s T-test is conducted via the functionality of the R stats package for every potential comparison between all groups in the given dataset. The resulting p-value for each comparison is displayed beneath the graph. If there is only one possible comparison (i.e., there are only 2 experimental groups), then only one p-value is shown; if more than one comparison is possible (i.e., there are 3 or more experimental groups), the p-values are presented in a grid format, where one group is identified on the y-axis and the group desired for comparison is listed on the x-axis, and then the corresponding p-value for the comparison can be viewed at the intersection of the grid coordinates for those groups. Datasets with only one sample per group do not have associated statistical comparisons.

**Figure 3.**
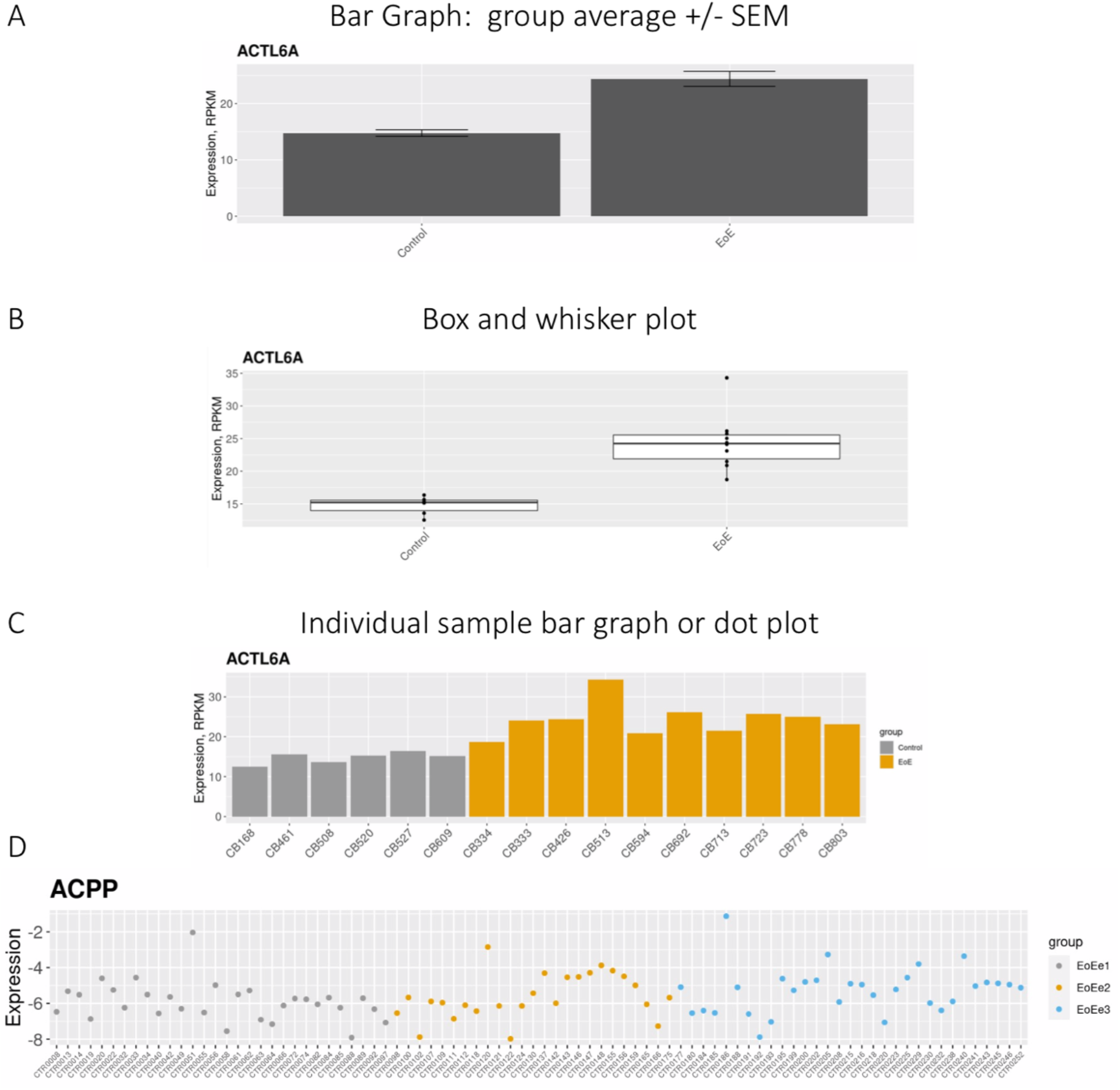
EGIDExpress graphical outputs. After choosing an experiment to browse from any category except single-cell RNAseq, the user chooses a graph output type. For all experiment types except single cell RNAseq, the choices are Bar graph: group average +/-SEM, Box and whisker plot, and either individual sample bar graph or dot plot. If the Bar graph option is chosen, a bar graph showing the mean of each experimental group is rendered for the given gene of interest, and error bars show the standard error of the mean (SEM). An example of such a graph is shown in (A). If the Box and whisker plot option is chosen, a box and whisker plot is rendered for the given gene of interest. The white box shows the interquartile range, the horizontal line within each box shows the median value, and the value for every individual sample is viewed as a dot. An example of such a plot is shown in (B). If one of the individual sample options is chosen, the data for a given gene are rendered either as a bar graph (e.g., [C]) or a dot plot (e.g., D).

In the first category of experiments, choosing a gene also renders data only for that gene in a tabular format in the second tab (Expression data). The information for this gene is filtered from the expression matrix and is made visible as a table in the second tab. This table lists the sample name in the first column, which matches the individual sample designations on the graphs, and the corresponding numeric expression value per sample is listed in the second column, the header of which specifies the unit of measurement.

In the first category of datasets, choosing a dataset displays its metadata and allows the user to download the expression matrix in the third and fourth tabs, respectively. The metadata is collected in table formatted file separate from the expression matrix; this is then imported into the RData file as an individual dataframe. In the third tab, Sample Information, a table is shown that lists the sample identifier and then also lists in separate columns relevant information about the sample such demographic, clinical, and histological information if the samples are derived from human subjects; sample grouping information; treatment information. The information is provided in a way that it can be linked to the identifiers provided in the graph and expression data tabs. In the fourth Download data, a download button in the tab allows for downloading a Unix/Linux CSV file of the expression matrix. Overall, any user can download the data and connect it to the information provided in the Sample Information tab to perform independent analysis.

The second category of datasets includes those in which the input is a Seurat object in RDS format from single-cell sequencing. A Seurat object is a collection of various data related to RNA-seq analysis with the Seurat package including expression data of single cells and metadata about each cell. Each dataset in this category has the option of up to three formats that are specified just under the gene dropdown menu and displayed in the Graph tab. The first possible option is Feature Plot (Fig. 4A). Choosing this option renders separate a UMAP plot for samples of each defined sample conditions, where the conditions are listed as the title of each graph. The gene name is listed at the right of all the maps. The x-axis is UMAP_1 and the y-axis is UMAP_2. The sum of all the cells from the given disease state are plotted on the UMAP plot corresponding to the appropriate sample conditions. Each dot represents one cell, and the intensity of the purple color is proportional to the expression level of that gene in the given cell. The names of the defined clusters are written in proximity to the given cluster. The second possible option is Violin Plot (Fig. 4B). For this option, a row that graphs all samples or multiple rows where samples are divided by disease state or other sample characteristic are shown. The y-axis shows the expression level, and the x-axis is divided by cell cluster, with the identity of the cluster identified by a key at the right. Each dot represents a cell. The third possible option is Ridge Plot (Fig. 4C). For this option, a row that graphs all samples or multiple rows where samples are divided by disease state or other sample characteristic are shown. The cluster identity is shown on the y-axis and each cluster is shown as an individual horizontal line on the graph/tiled, with the identity of the cluster indicated by the key to the right of the graph. The expression level is indicated on the x-axis, ascending from left to right. The height of the graph for a given cluster at a given x-axis coordinate is proportional to the number of cells that exhibit expression at that particular value. For some datasets, such as the T cell single cell RNAseq, a checkbox exists just below the option for the graph types that allows for the data to be visualized with all cells in the same graph or divided by disease state/sample characteristics.

**Figure 4.**
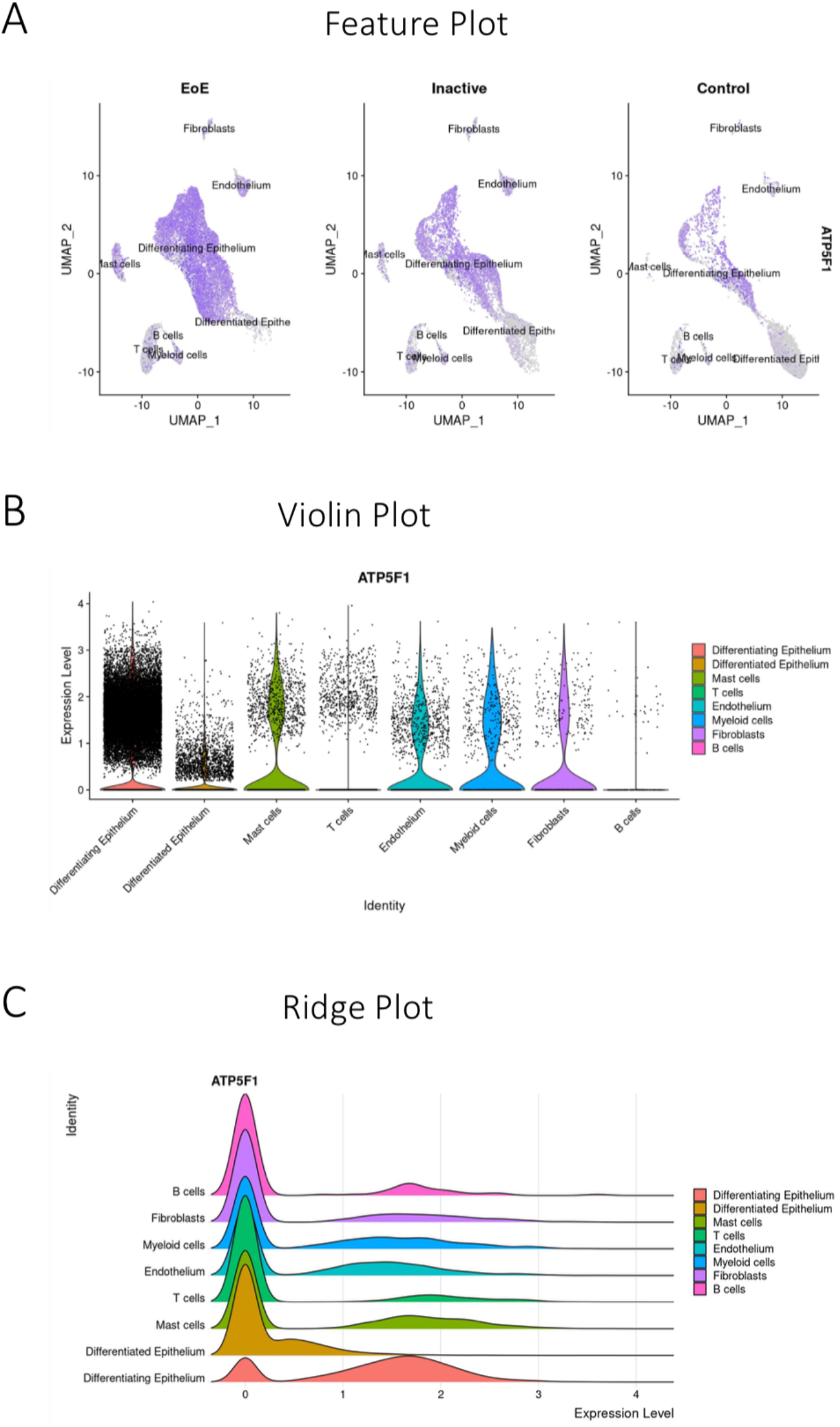
EGIDExpress single-cell RNAseq graphical outputs. If the user chooses to browse an experiment in the single-cell RNAseq category, the user can choose one of two graph types to visualize the data for the chosen gene of interest, namely a feature plot (A), a violin plot (B) or a ridge plot (C).

The third category of datasets, CHIPseq, uses BAM files as inputs. These files are binary files of sequence alignment data derived from FASTQ files. EGIDExpress generates a visualization in a genome browser style, where the strength of signal for a given chromatin modification at a given genomic coordinate among samples in a given experiment is graphed. At the top of the graph, the chromosome on which the given gene is located is shown, along with its banding pattern (Fig. 5). A red slider box indicates the region of the chromosome that is visualized in the tracks below it. A track just below the chromosome illustration shows known transcripts in the region (indicated by horizontal lines), with exons indicated by vertical lines. The scale for this track is indicated at the top, and this scale also applies to all the tracks beneath it. For each chromatin modification, the resulting graph’s x-axis indicates the chromosomal coordinate, and the y-axis indicates the strength of signal for the given chromatin modification at that coordinate. Islands that are statistically significant previously identified by MaNORM for each given chromatin modification and are indicated by a horizontal bar below the chromosomal coordinates that make up the island.

**Figure 5.**
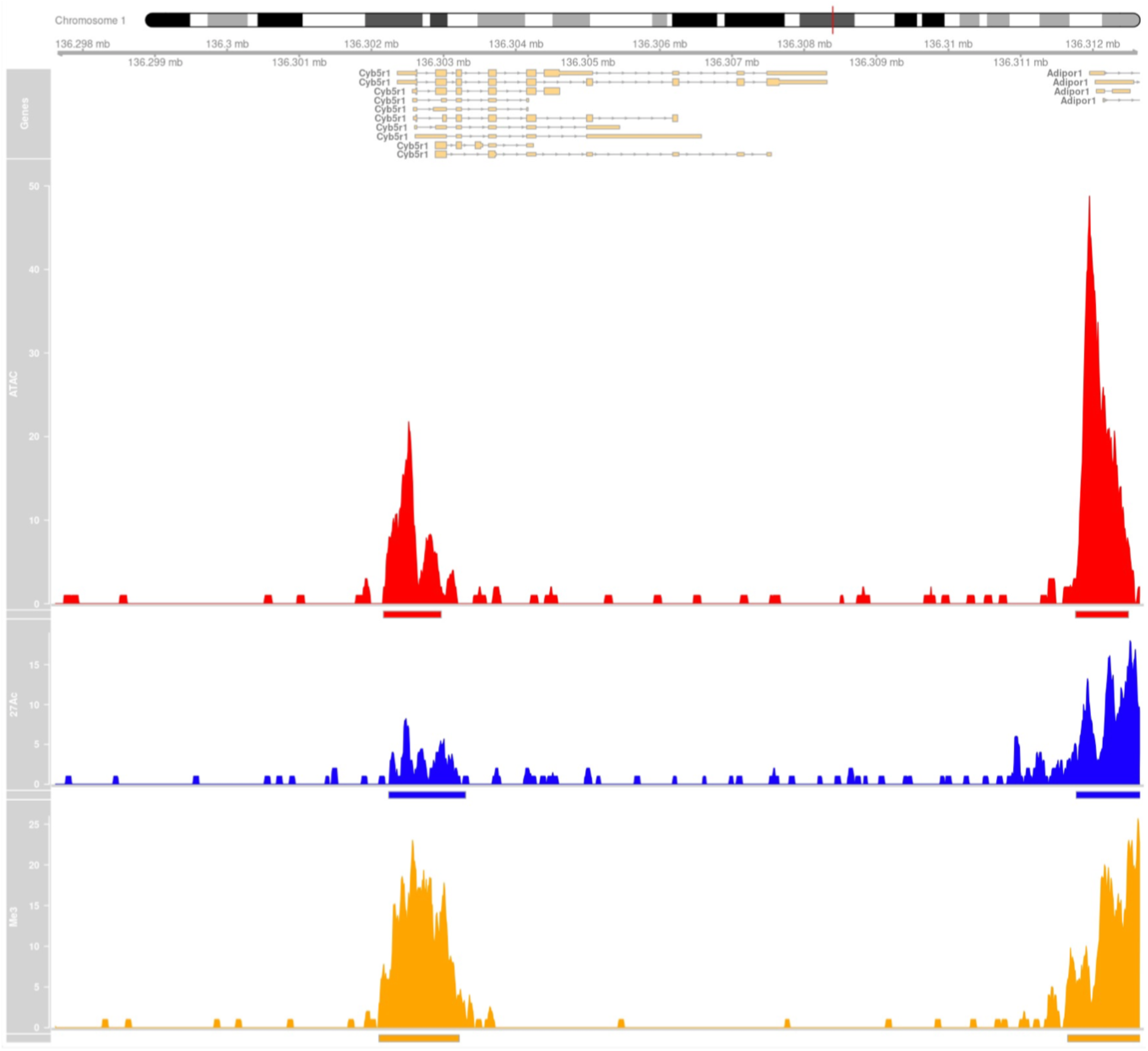
EGIDExpress ChIPseq graphical outputs. If the user chooses to browse an experiment in the ChIPseq category, the graphical output shows the chromosome on which the given gene is located (top) and four additional tracks: first, a track is shown with the scale of the local chromosomal environment as specified by the user (number of basepairs relative to the transcriptional start site for the given gene). Next, all known transcripts located within that region are shown. The bottom three tracks show data for chromatin modifications assessed within the chosen experiment. Here, results from ATACseq (red), histone H3 K27 acetylation (blue), and H3K4 tri-methylation (yellow) are shown. For each track, islands that are statistically significant are indicated by bars of the same color as the graphs underneath the peaks.

## Results and Discussion

One of the goals in developing EGIDExpress included production of a platform that is easily amenable to adding, distributing, and visually sharing data with colleagues and collaborators. As of September 2022, there are 19 datasets on the public site and 18 datasets on the private site. To date (2.8 years in existence), over 2,400 users have visited the sites, including from 37 countries and from 37 states within the USA. Several features of the design and implementation of EGIDExpress allow us to achieve and refine our work toward these goals.

The current design of EGIDExpress allows for convenient addition of several types of datasets, including those that can be expressed in the form of an expression matrix, RDS files, or BAM files. The current architecture allows even the novice to be able to add datasets to the website with the use of job aides. Upon soliciting expression matrix, RDS, or BAM files and associated information, the time to add it to the website is minimal (less than 30 minutes). One challenge is making sure that the associated information is provided so that the dataset is complete. Many types of biological datasets can be fitted into one of these formats. This adds to the versatility of EGIDExpress.

EGIDExpress allows for easy distribution of data among many individuals. EGIDExpress is free and easily available to multiple users simultaneously. Compared to other publicly available repositories (e.g., NCBI GEO) or software (e.g., GeneSpring GX), we suggest it has a more intuitive, simple interface. This is especially true if a user wants to approach multiple datasets with the intent of checking the expression levels of only one or a limited number of genes. The simple interface also allows for quick understanding of the experiment and experimental groupings as well a link to the primary publication all on the same page. Of note, any datasets derived from human samples have been de-identified prior to their addition to the database. Despite these advantages, EGIDExpress does not yet house as many datasets as other public repositories.

As evidence for the ease of distribution of the data from EGIDExpress and visibility to the world at large, we have academic collaborators and pharmaceutical companies who have downloaded the data. A pharmaceutical company has already written an article based on the data derived EGIDExpress, proving that we are accelerating hypothesis generation and new data analysis ^13^. Deposition of a dataset on EGIDExpress has already in one case satisfied a journal’s requirement for data deposition in a publicly available repository ^14^. Published data are freely available for individuals who have personal interest in learning about EGIDs and the research occurring in the Rothenberg CURED Lab, such as patients or patient advocacy groups; they can browse the site for their own scientific efforts or to see the progress being made in the lab. To some degree, we can determine basic information about site users using google analytics; this allows us to infer if individuals at a given university or conference may be amenable to engaging in collaborations with us. The home page has information about the Rothenberg CURED Lab including contact information to encourage users of the website to contact us directly if they have questions, want to see our advise and/or mentorship in establishing a similar app for their own data, want to collaborate with us, or learn more about our laboratory.

In the academic research laboratory environment, EGIDExpress has the advantage of allowing data to be shared among multiple lab members. It promotes continuity of data transfer, data integrity, and prevents data loss as individuals leave the lab. EGIDExpress is a read-only interface; there is no way for users to edit data in EGIDExpress, so there is no risk of data loss or inappropriate modification. On the back end, EGIDExpress motivates individuals to deposit raw data files, sample information, and experiment information in a specified location so that future lab members can access all the information together in an interpretable way. This can save the future users time, promote data integrity, and extend data usage to projects not envisioned at the time the experiment was originally performed. Overall, to date EGIDExpress has shown that it has the potential to distribute data among local and external colleagues and collaborators in academia and industry.

The final goal of producing EGIDExpress was to facilitate visually sharing data among colleagues, collaborators, and other interested individuals. EGIDExpress allows for quick browsing of multiple datasets on a per gene basis. Datasets that are in the format of an expression matrix can be visualized in one of several different graph options, including bar graph, box and whiskers plot, and individual bar or dot graphs. Datasets that are in the format of an RDS file can be viewed as feature plots, violin plots, or ridge plots. Datasets that are in the format of BAM files can be viewed in a genome browser format.

Current laboratory members frequently utilize EGIDExpress to screen capture pictures of graphs for unofficial presentations such as internal laboratory meetings. The graphs produced by EGIDExpress can also be downloaded in various image formats or viewed in higher resolution by right clicking on the graph in any browser and clicking Save image as or Open image in new tab. Raw data are also readily available on EGIDExpress so that individuals can transfer them to other graphing or analysis programs. We suggest that the data are presented in a format that is immediately useful for collaborative science.

Some drawbacks of EGIDExpress include that the graphs are not as customizable as they are in other programs, and different visualization choices and graph types are relatively limited. However, the download data functionality provides a means to capture the input of the EGIDExpress analysis (the expression matrix) which can be put into a program such as GeneSpring if more complicated or customized analyses are desired. Another drawback is that in some cases, only data that have been processed or normalized to some degree are presented on EGIDExpress, and the raw data for experiments are not available to users via the website (e.g., FASTQ files for RNAseq experiments, or RDS files for single cell experiments). The data visualized in some cases thus depends on QC/pre-processing/normalization steps; for example, in some cases minimum expression cutoffs may impact the genes that are included in the dataset on the website. However, the raw data files for all experiments are kept on the back end in an easily accessible, organized structure so that they can be accessed and potentially available to collaborators upon request. Graphs and statistics presented on EGIDExpress are not downloadable in bulk; the user must browse genes individually to observe the graphs and/or statistics. Another disadvantage of EGIDExpress is that the names of the same gene may differ among various datasets in EGIDExpress, making computational direct comparisons among datasets challenging. One challenge related to implementation of EGIDExpress involves ensuring that the server has the most updated version of R and all appropriate packages installed. Despite these challenges and drawbacks, we will continue to strive to improve EGIDExpress to optimize its utility and reach.

EGIDExpress aims to contribute to understanding the molecular basis for EGIDs, which is an important scientific and health problem. These diseases affect many people worldwide, causing them to suffer a low quality of life. At the same time, these are rare diseases, especially EGIDs that affect the stomach, small intestine, and colon. As such, research is lacking and relevant biological specimens for study are scarce. By making data publicly available to researchers worldwide who are addressing this problem, we hope to accelerate identification of diagnostics, treatments, and ultimately cure for the disease. The data shared are from patient-derived samples from individuals with rare diseases, so sharing as much data with as many researchers as possible within and across disciplines may help promote advances that were not previously possible. By reaching a wider audience, people with novel thoughts and ideas from different fields have approached us to form collaborations and to share novel insights beyond what our research team had alone. Overall, through EGIDExpress, we hope to maximize the utility and usage of data so that we and others can make progress toward the development of life-improving interventions that will impact the lives of people with these diseases.

In summary, we have developed a user-friendly web app that allows for the easy addition and wide distribution of large, published datasets to the public at large and for distribution of unpublished datasets among colleagues. Multiple types of datasets can be quickly added to the website. A wide range of users can browse the website or download graphs or datasets for visualization or further analysis. Since its inception in 2019, it has demonstrated its utility at the local laboratory level and beyond such as by generating interest from collaborators in academia and industry. At least 12 publications have already been facilitated by EGIDExpress ^15, 16, 17, 18, 19,20, 21, 14, 22, 23, 24, 25^. Datasets will continue to be added to EGIDExpress as they are generated, and we remain flexible to adding new data formats or modes of visualization as new dataset types arise, and we are open to developing new functionalities within the app. Overall, we will continue to pursue the goals of EGIDExpress to accelerate research toward finding better diagnostics, treatments, and a cure for EGIDs.

